# Skin epithelial cells change their mechanics and proliferation upon Snail-mediated EMT signalling

**DOI:** 10.1101/2022.02.01.478626

**Authors:** Kamran Hosseini, Palina Trus, Annika Frenzel, Carsten Werner, Elisabeth Fischer-Friedrich

## Abstract

Skin cancer is the most commonly occurring cancer in the USA and Germany, and the fourth most common cancer worldwide. Snail-dependent epithelial-mesenchymal transition (EMT) was shown to initiate and promote skin cancer. Previous studies could show that EMT changes actin cortex regulation and cellular mechanics in epithelial cells of diverse tissue origin. However, in spite of its potentially high significance in the context of skin cancer, the effect of EMT on cellular mechanics, mitotic rounding and proliferation has not been studied in skin epithelial cells so far. In this work we show that TGF-*β*-induced partial EMT results in a transformation of the mechanical phenotype of skin epithelial cells in a cell-cycle dependent manner. Concomitantly, we looked at EMT-induced changes of cell proliferation. While EMT decreases proliferation in 2D culture, we observed an EMT-induced boost of cellular proliferation when culturing cells as mechanically confined aggregates of skin epithelial cells. This proliferation boost was accompanied by enhanced mitotic rounding and composition changes of the actin cortex. We give evidence that observed EMT-induced changes depend on the EMT-upregulated transcription factor Snail. Overall, our findings indicate that EMT-induced changes of cellular mechanics might play a currently unappreciated role in EMT-induced promotion of skin tumor proliferation.

**Significance statement:** This study describes how epithelial-mesenchymal transition (EMT) alters the actin cytoskeleton, cellular mechanics and proliferation in a benign tumor model of skin epithelial cells. We show that corresponding EMT-induced phenotypes depend on the signalling of the transcription factor Snail. Our findings suggest that EMT-induced changes of cellular mechanics and proliferation might play a currently under-appreciated role in EMT-induced promotion of skin tumors.

## 1. Introduction

Skin cancer has been reported as the most commonly occurring cancer in the USA and Germany [1, 2] and the fourth most common cancer worldwide [3]. Furthermore, instigation and promotion of skin cancer has been shown to be driven by epithelial-mesenchymal transition (EMT) [4].

EMT is a cellular transformation that results in epithelial cells losing their epithelial-like properties and gaining mesenchymal-like properties. In particular, typical EMT-induced changes include the loss of apical-basal polarity and cell-cell adhesion, the adoption of a spindle shaped morphology, increased cell motility as well as characteristic changes in the cytoskeleton [5–7]. If cells undergo EMT in an incomplete manner, i.e. if they transition from an epithelial state to a state with mixed epithelial and mesenchymal traits, the transformation is called partial EMT [8]. In tumor progression, EMT and partial EMT are known to make cancer cells more invasive [5–7, 9, 10] and more or less proliferative depending on their microenvironment and their malignancy level [11–13].

EMT signalling is mediated by several transcription factors in the cell including Snail [14, 15]. Snail activation was shown to induce EMT in cancer and embryonic development [14, 15]. Furthermore, Snail was shown to drive cancer initiation and progression [4]. We could previously show that EMT promotes cell proliferation and mitotic rounding in confined tumor spheroids of breast cancer cells [11]. Changes in proliferation and mitotic cell morphology were accompanied by changes in mechanics and composition of the actin cortex [11, 16]. Despite the importance of EMT in the context of skin cancer, there are so far no studies on the effect of EMT on cellular mechanics and proliferation in skin cancer models. To further advance the knowledge in this field, we present here a study on the effects of TGF-*β*-induced partial EMT on cell mechanics and proliferation on HaCaT skin epithelial cells.

HaCaT cells are a cell line of human keratinocytes that spontaneously immortalized. This immortalization could, at least in part, be traced back to mutations in the tumor suppressor gene p53, whose mutation spectrum is typical for ultraviolet light induced mutations [17]. Correspondingly, HaCaT cells constitute a model for benign skin cancer [18]. Next to investigating cortex-mechanical properties of isolated HaCaT cells with and without EMT-induction, we monitored the effect of EMT on HaCaT spheroid growth and mitotic roundness. We further investigated the effect of the transcription factor Snail on actin cortex mechanics and cell proliferation.

## 2. Results and discussion

We induced EMT signalling in HaCaT skin epithelial cells, through an established method by incubation with TGF-*β* 1 [19, 20], (Figure S1, Supplementary information and Experimental section). In accordance with typical EMT proteomic changes, HaCaT cells showed a treatment-induced reduction of the epithelial marker E-cadherin and an increase in mesenchymal markers vimentin and N-cadherin according to Western blot quantification and immunofluorescence (Figure S1b-c,f, Supplementary information). Partial retention of epithelial traits such as moderate cell-cell adhesion and E-cadherin expression is in accordance with earlier reports that this treatment leads to a partial rather than a complete EMT [20]. TGF-*β*1 treatment enhances migration of HaCaT cells corresponding to a mesenchymal cell phenotype, while reducing its proliferation in 2D culture (see [19] and Figure S1d, Supplementary information). In the following, we will denote HaCaT cells that underwent partial EMT as modHaCaT cells.

### 2.1. EMT affects cortex mechanics and composition in skin epithelial cells

Motivated by previous observations of EMT-induced changes of actin cortex mechanics in breast, lung and prostate epithelial cells [11, 16], we wanted to test if EMT also changes the mechanics of the actin cortex in HaCaT skin epithelial cells. For this purpose, we measured untreated and EMT-induced cells with a cell-mechanical assay based on dynamic cell confinement with the cantilever of an atomic force microscope (AFM) (Figure 1a-b, see Experimental section). This assay has been previously established as a method for the measurement of material properties of the actin cortex [11, 16, 21–25]. In particular, it provides for each measured cell a readout of i) cortical tension, ii) cortical stiffness and iii) phase shift at a specific frequency of applied deformations. (While cortical stiffness rates the overall mechanical resistance of the material towards deformation, the phase shift reports the viscoelastic nature of the material and adopts a value between 0° and 90°, where 0° corresponds to an entirely solid-like and 90° corresponds to an entirely fluid-like material.)

**Figure 1.**
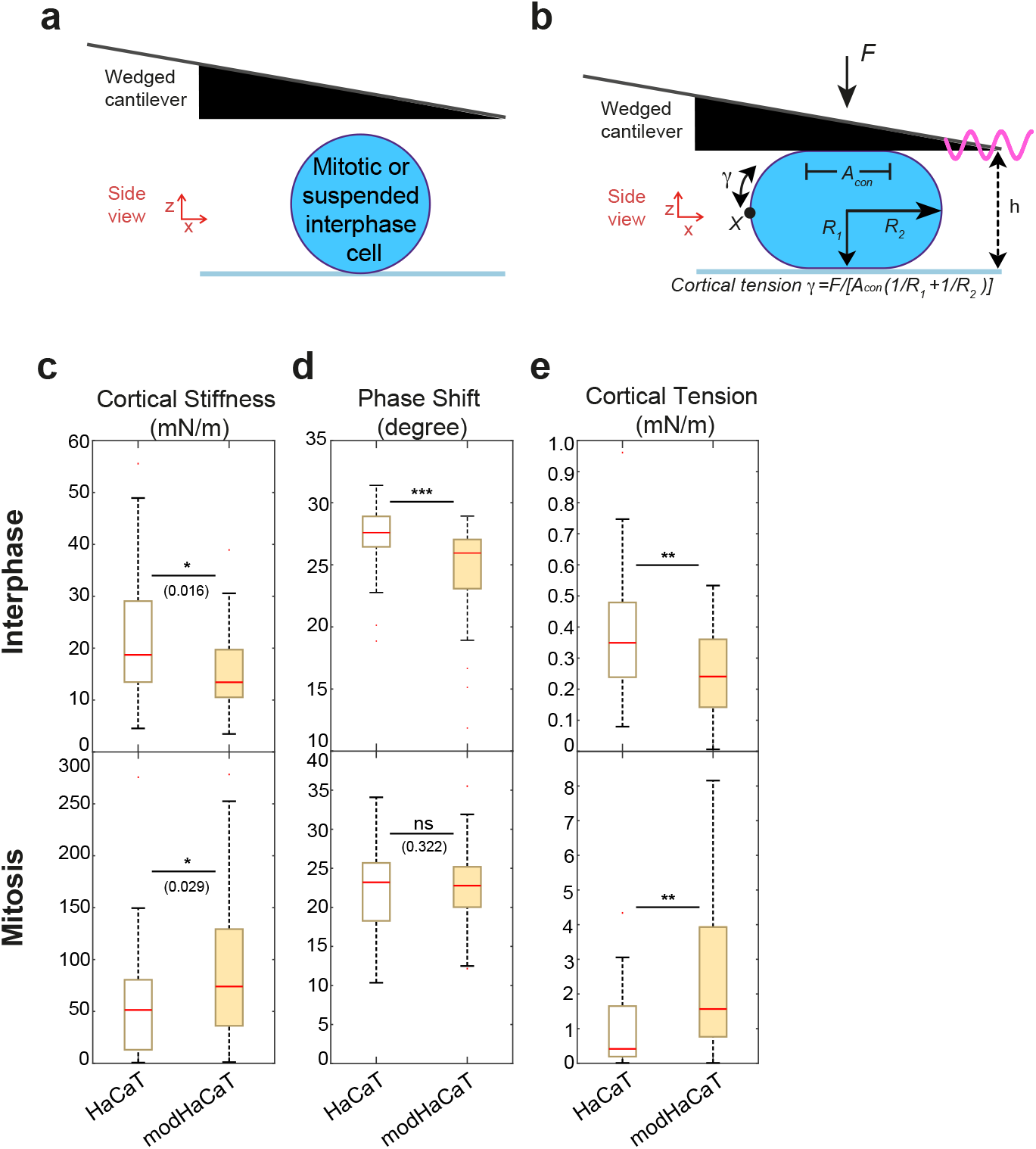
Actin cortex mechanics of HaCaT skin epithelial cells before and after EMT measured by dynamic AFM cell confinement. a,b) Schematic of measurement setup: initially round cells (mitotic cells or suspended interphase cells) are confined by the tip of a wedged AFM cantilever and oscillatory height oscillations are imposed. b) Geometrical parameters of a cell estimated during AFM confinement. c) Cortical stiffness, d) phase shift, and e) cortical tension measured for suspended interphase cells (top row) and STC-arrested cells in mitosis (bottom row) before and after EMT. Post-EMT cells are referred to as modHaCaT. Yellow-shaded boxes indicate post-EMT conditions. Number of cells measured: Interphase: HaCaT n = 36, modHaCaT n = 29, Mitosis: HaCaT n = 20, modHaCaT n = 17. Measurements are representative for two independent experiments. n.s.: *p* > 0.05, * : *p* < 0.05, ** : *p* < 0.01, *** : *p* < 0.001.

Our cell-mechanical assay relies on an initially spherical cell shape. Hence, interphase cells were measured in suspension. This setup choice also increases comparability of measurements of interphase cells versus rounded mitotic cells. Our measurements on interphase cells showed an EMT-induced decrease in cortical tension and stiffness (Figure 1c,e, upper row). In addition, we find that the measured phase shift is decreased in modHaCaT interphase cells indicating a solidification of the cortical material (Figure 1d, upper row). Increased cortical solidification in conjunction with stiffness reduction has been shown before in response to myosin-II inhibition [21].

Subsequently, we assessed the mechanical properties of the cell cortex in mitotic cells before and after EMT. For mechanical measurements, cells were pharmacologically arrested in mitosis by addition of S-trityl-L-cysteine (STC) to enrich mitotic cells and prevent cell-cycle-related mechanical changes (see Experimental section) [11, 16, 23, 24, 26]. STC-treated cells were previously shown to exhibit cell-mechanical properties comparable to cells in metaphase of mitosis [11, 23]. We saw a mechanical change in mitosis opposite to changes in interphase HaCaT cells: cortical tension and cortical stiffness were clearly increased in HaCaT cells while phase shifts remained unchanged (Figure 1c-e, lower row). This observation is in accordance with observations for other epithelial cell lines from different tissue origin [11, 16]. In short, our data shows that the EMT-induced mitotic cortex has a higher stiffness and contractility.

In order to better understand cell-mechanical changes upon EMT on a molecular level, we investigated EMT-induced changes of actin and myosin-II levels in HaCaT cells. First, we quantified total levels of actin and myosin-II by western blotting before and after EMT in asynchronous cell populations (mainly interphase) and in mitotic cells (Figure 2a). Analogous to previous observations in breast epithelial cells [11], we observed no significant changes in expression levels of asynchronous cell populations (mainly interphase cells). For mitotic cells, we found an EMT-induced increase of myosin-II (Figure 2a), while there were no changes of actin abundance.

**Figure 2.**
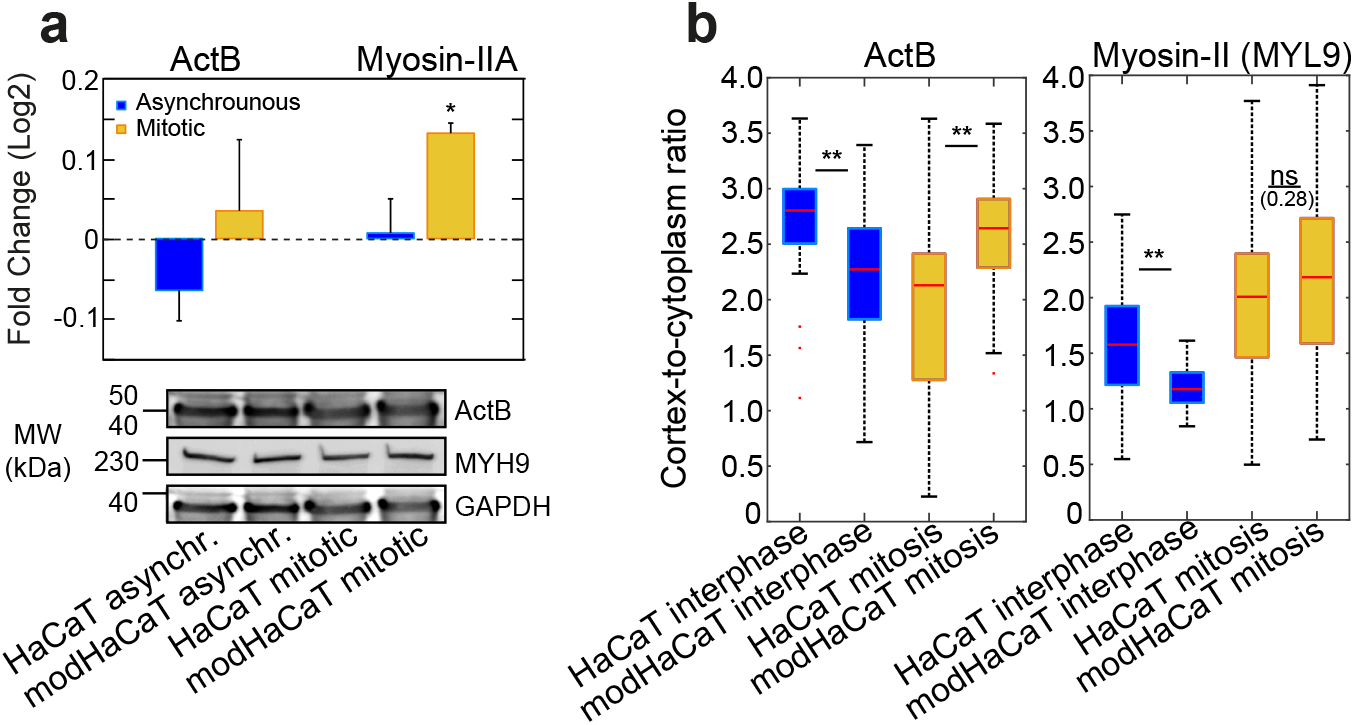
Changes of actin and myosin-II abundance and distribution upon EMT in HaCaT skin epithelial cells. a)Relative changes of actin and myosin-II expression from Western blots. Top panel: Quantification of relative changes (modHa-CaT/HaCaT). Quantification was done against GAPDH. Bottom panel: Western blots showing actin (ActB) (top row) and myosin-IIA (MYH9) (second row) abundance before and after EMT in asynchronous cell populations (mainly interphase) and mitotic cells. b) Ratio of cortical versus cytoplasmic actin (ActB) (left panel) and myosin-II (MYL9) (right panel) in suspended interphase cells and STC-arrested mitotic cells upon EMT. Post-EMT cells are referred to as modHaCaT. Blue-shaded boxes indicate interphase cells while yellow-shaded boxes indicate mitotic cells. Number of samples measured in (a): n=2; in (b): ActB: HaCaT interphase n=31, modHaCaT interphase n=32, HaCaT mitotic n=30, modHaCaT mitotic n=30. Myosin-II (MYL9): HaCaT interphase n=31, modHaCaT interphase n=32, HaCaT mitotic n=32, modHaCaT mitotic n=31. Measurements are representative for at least two independent experiments. n.s.: *p* > 0.05, * : *p* < 0.05, ** : *p* < 0.01, *** : *p* < 0.001.

In another set of experiments, we quantified the cortex-to-cytoplasm ratio of actin and myosin-II before and after EMT (Figure 2b). To this end, we transfected HaCaT cells before and after EMT with plasmids expressing fluorescently labeled actin (ACTB-mCherry) or myosin light chain 9 (Myl9-mApple) (see Experimental section). Subsequently, we imaged the equatorial cross-section of these cells and quantified the ratio of cortical versus cytoplasmic fluorescence intensity (Figure 2b and Figure S2a-d, Supplementary information, see Experimental section). We found that the cortex-to-cytoplasm ratio of actin and myosin-II decreases in interphase cells upon EMT (Figure 2b, blue-shaded boxes). By contrast, in mitosis, we observed an increase of the actin cortex-to-cytoplasm ratio, but no change of the myosin-II cortex-to-cytoplasm ratio (Figure 2b, yellow-shaded boxes). These changes of cortex-to-cytoplasm ratio are qualitatively similar to previously quantified cortex-to-cytoplasm ratios for breast epithelial cells [11]. However, in breast epithelial cells, we did not observe a decrease of actin cortex-to-cytoplasm ratio upon EMT in interphase cells. Considering both experimental findings together (western blotting and cortex-to-cytoplasm ratios), we conclude that the amount of cortex-associated actin increases in mitosis but decreases in interphase cells. In addition, cortical myosin-II decreases in interphase upon EMT and overall abundance of myosin-II increases in mitosis (Figure 2). EMT-induced changes of actin and myosin-II may thus provide a simple mechanistic explanation for EMT-induced cortex-mechanical trend.

Previously, it has been shown that also the intermediate filament protein vimentin plays a major role in the generation of cortical tension in mitosis [27]. To investigate the contribution of vimentin to cortical tension changes upon EMT, we performed a knock-down of vimentin by RNA interference (RNAi) (Figure S4, Supplementary information). We then applied our mechanical assay on STC-arrested mitotic HaCaT and modHaCaT cells with and without knockdown of vimentin (Figure S4a-c, Supplementary information). We observed a reduction of cortical tension and stiffness in both pre- and post-EMT HaCaT cells upon vimentin knock-down corroborating the role of vimentin for cortical mechanics in mitosis (Figure S4a-c, Supplementary information). Cortical tension and stiffness decrease were larger in post-EMT cells suggesting that EMT-induced increase of cortical vimentin makes a contribution to the EMT-induced mitotic phenotype of cortical mechanics in addition to EMT-induced changes of actin and myosin (Figure S4a,c, Supplementary information).

### 2.2. Snail affects cortical mechanics in a cell-cycle dependent manner

The transcription factor Snail regulates induction of EMT in cells e.g. during embryonic development [15]. In addition, Snail has been shown to promote cancer likely through promotion of EMT [4]. Consequently, we checked the expression of Snail changes upon EMT in HaCaT skin epithelial cells. In accordance with previous results, we observed a significant EMT-induced increase of Snail expression (Figure S1g and S3a-b, Supplementary information) [6, 28, 29].

In order to test the cell-mechanical effect of Snail in HaCaT skin epithelial cells, we performed a knock-down of SNAI1 by RNA interference (RNAi), leading to a decreased expression of Snail (Figure S3a-b, Supplementary information). We then performed our mechanical assay in interphase and mitotic HaCaT and modHaCaT cells with and without knock-down of SNAI1 (Figure 3). We verified that knock-down of SNAI1 results in significant reduction of overall Snail protein in post-EMT cells (Figure S3a-b), indicating that SNAI1 is the dominant Snail isoform in post-EMT cells.

**Figure 3.**
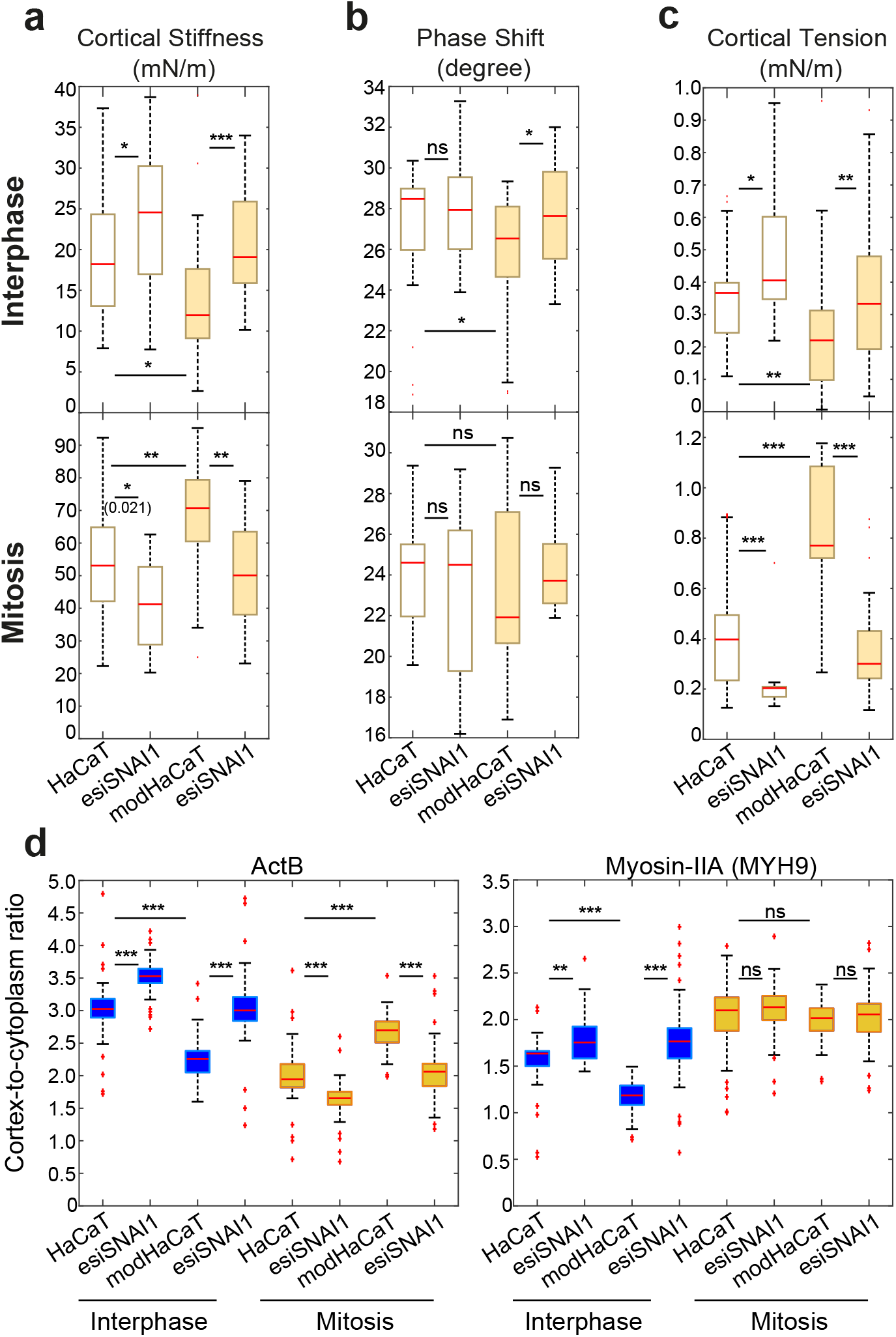
Effect of SNAI1 knock-down on cortex mechanics in HaCaT skin epithelial cells with and without EMT. a) Cortical stiffness, b) phase shift, and c) cortical tension measured by dynamic AFM confinement. Top row: suspended interphase cells. Bottom row: STC-arrested mitotic cells. d) Ratio of cortical versus cytoplasmic actin (ActB) and myosin-IIA (MYH9) in suspended interphase and STC-arrested mitotic HaCaT cells upon EMT and SNAI1 knock-down. Post-EMT HaCaT cells are referred to as modHaCaT. Yellow-shaded boxes in (a-c) indicate post-EMT conditions. Blue-shaded boxes indicate interphase cells while yellow-shaded boxes indicate mitotic cells in (d). Number of cells measured in (a-c): Interphase: HaCaT n = 27, esiSNAIl n = 28, modHaCaT n = 27, esiSNAIl n = 29, Mitosis: HaCaT n = 22, esiSNAIl n = 20, modHaCaT n = 20, esiSNAI1 n = 19. (d): ActB: Interphase: HaCaT n = 36, esiSNAIl n = 39, modHaCaT n = 31, esiSNAIl n = 36, Mitosis: HaCaT n = 34, esiSNAIl n = 38, modHaCaT n = 35, esiSNAI1 n = 34. Myosin-IIA (MYH9): Interphase: HaCaT n = 36, esiSNAIl n = 36, modHaCaT n = 35, esiSNAIl n = 37, Mitosis: HaCaT n = 32, esiSNAIl n = 39, modHaCaT n = 28, esiSNAI1 n = 37. Measurements are representative for at least two independent experiments. n.s.: *p* > 0.05, * : *p* < 0.05, ** : *p* < 0.01, *** : *p* < 0.001.

In non-transformed interphase cells (HaCaT), we observed that cortical tension and elastic modulus increase after SNAI1 knock-down, while phase shift does not show significant changes (Figure 3a-c, upper row). Vice versa, we observed a decrease of cortical tension and elastic modulus in mitotic cells, while phase shift remained unchanged (Figure 3a-c, lower row). In post-EMT cells (modHaCaT), cortical changes had the same trend but changes were stronger compared to pre-EMT cells (Figure 3a-c, yellow-shaded boxes).

Overall, we conclude that Snail signalling affects actin cortex mechanics, and its knock-down causes changes of cortical tension and stiffness. Corresponding cell-mechanical changes are opposite in interphase and mitosis. Furthermore, SNAI1 knock-down restores the epithelial cell-mechanical phenotype in post-EMT cells (Figure 3, yellow-shaded boxes).

Concomitantly, we quantified the cortex-to-cytoplasm ratio of actin and myosin-II in suspended interphase and STC-arrested mitotic HaCaT cells before and after EMT with and without knock-down of SNAI1 (Figure 3d, see Experimental section). We see that SNAI1 knock-down leads to an increased cortex-to-cytoplasm ratio of actin in suspended interphase HaCaT cells (Figure 3d, blue-shaded boxes). On the other hand, in mitosis, SNAI1 knock-down leads to a decreased cortex-to-cytoplasm ratio of actin (Figure 3d, yellow-shaded boxes). Therefore, SNAI1 knock-down induces changes of cortical actin that are opposite to EMT-induced changes. For myosin-II, we observed an increase of the cortex-to-cytoplasm ratio upon SNAI1 knock-down in interphase cells (Figure 3d, yellow-shaded boxes). Again, this myosin-II phenotype was opposite to EMT-induced changes. In mitosis, the cortex-to-cytoplasm ratio of myosin-II is not affected by SNAI1 knock-down. In conclusion, we found that the cortical levels of actin and myosin-II in post-EMT HaCaT cells are qualitatively restored to pre-EMT values in both interphase and mitosis upon SNAI1 knock-down (Figure 3d).

### 2.3. Mitotic rounding and proliferation are enhanced in confined spheroids upon EMT

Mitotic rounding is a cellular shape change at the onset of mitosis that is widely conserved amongst animal cells [30]. It describes the process of cells leaving their interphase shape and adopting an almost spherical shape at mitotic onset. Through mitotic rounding, a well-defined space for the formation of the mitotic spindle is provided in the cell interior. It was shown previously that mitotic rounding is a prerequisite for timely and successful cell division [31–34]. As opposed to cells growing in 2D culture, (tumor) cells in the body are mechanically confined by other cells or stroma. This confinement poses a mechanical hindrance towards cell shape changes such as mitotic rounding. This hindrance needs to be overcome by cellular force generation. It was previously shown that a major driving force for mitotic rounding is actin cortex contractility [11, 22, 23, 26, 35–38]. Therefore, our finding of increased cortical tension in mitosis suggests that mitotic rounding might be enhanced in HaCaT skin epithelial cells.

We thus investigated whether EMT-induced changes of the actin cortex also changes mitotic rounding of HaCaT cells in a mechanically confining environment. Furthermore, we checked if those changes in mitotic rounding correlate with a change in cell proliferation.

For this purpose, we seeded skin epithelial cells in 3D culture embedded in elastic non-degradable poly(ethylene glycol) (PEG)-heparin gel. Over time, individual suspended cells grew into spheroidal cell aggregates which were mechanically confined by the gel. After one week of growth (including the pharmacological treatment time for EMT induction, before being fixed and immunostained), spheroids were fixed and stained (see Experimental section). Then, spheroids were imaged through confocal z-stacks. Thereafter, we quantified the growth of spheroid size by measuring the largest cross-sectional area of the spheroid and the corresponding number of cells in that cross-section (Figure 4b-c). Furthermore, we quantified the roundness of mitotic cells in fixed spheroids with and without EMT induction (Figure 4d). STC (2*μ*M) was added to arrest cells in mitosis 24 hours before fixation in order to enrich mitotic cells in spheroids. For each mitotic cell, the largest cross-sectional area of the cell was used to quantify roundness (Figure 4a, middle panels, see Experimental section).

**Figure 4.**
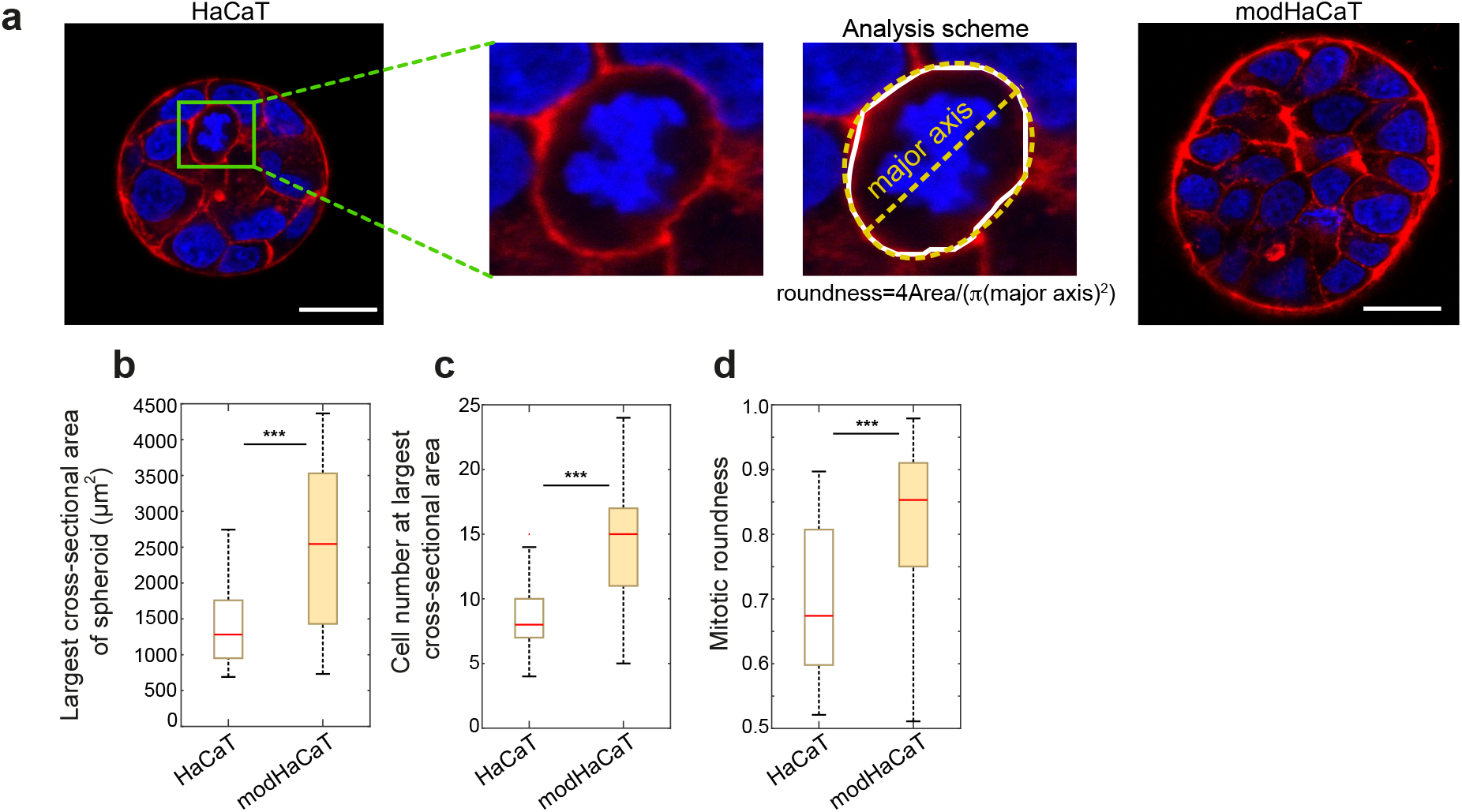
EMT enhances proliferation and mitotic roundness in HaCaT spheroids. a) Outer panels: Confocal images of cross-sections of HaCaT and modHaCaT spheroids in elastic PEG-heparin gels (Young’s modulus of ≈3 kPa) after 1 week of growth. Cells were fixed and stained for DNA (DAPI, blue) and F-actin (Phalloidin, red). Mitotic cells were arrested in mitosis through co-incubation with STC (2 *μ*M) prior to fixation and exhibit condensed chromosomes (see e.g. cells in green frame). Scale bar: 20 *μ*m. Middle panels: zoom of cross-sections of mitotic cells. For image analysis, the cell outline of the largest cell cross-section was determined (white line) and then an ellipse was fitted to determine roundness, see Experimental section. b-d) Spheroid size, cell number and mitotic roundness with and without EMT. b) Spheroid size was quantified by the largest cross-sectional area of the spheroid. c) Cell number was quantified as the number of cells in the maximum cross-sectional area of the spheroid (see Experimental section). Post-EMT HaCaT cells are referred to as modHaCaT. Yellow-shaded boxes indicate post-EMT conditions. Number of spheroids analyzed: (b-c): HaCaT n = 35, modHaCaT n = 35. Number of mitotic cells analyzed (d): HaCaT n = 29, modHaCaT, n = 45.Measurements are representative for at least two independent experiments. n.s.: *p* > 0.05, * : *p* < 0.05, ** : *p* < 0.01, * * * : *p* < 0.001.

Our analysis shows an increased proliferation and mitotic roundness upon EMT (Figure 4b-d). In fact, the proliferation trend in this mechanically confining environment is opposite to the 2D culture, where EMT leads to reduced proliferation (Figure S1d, Supplementary information).

These results suggest that EMT-induced cell-mechanical changes enhance mitotic rounding of HaCaT skin epithelial cells. We speculate that this, in turn, could contribute to enhanced proliferation e.g. through reduced mitotic duration [31, 39, 40].

### 2.4. Snail knock-down rescues EMT-associated phenotypes in confined spheroids

To investigate the effect of Snail expression changes on HaCaT skin epithelial cells in 3D culture, we performed an RNAi knock-down of SNAI1 in spheroids (see Figure 5a, and Experimental section). SNAI1 knock-down led to a significant decrease of spheroid proliferation as well as mitotic roundness of cells (Figure 5b-d), more so for post-EMT cells (Figure 5b-d, yellow-shaded boxes).

**Figure 5.**
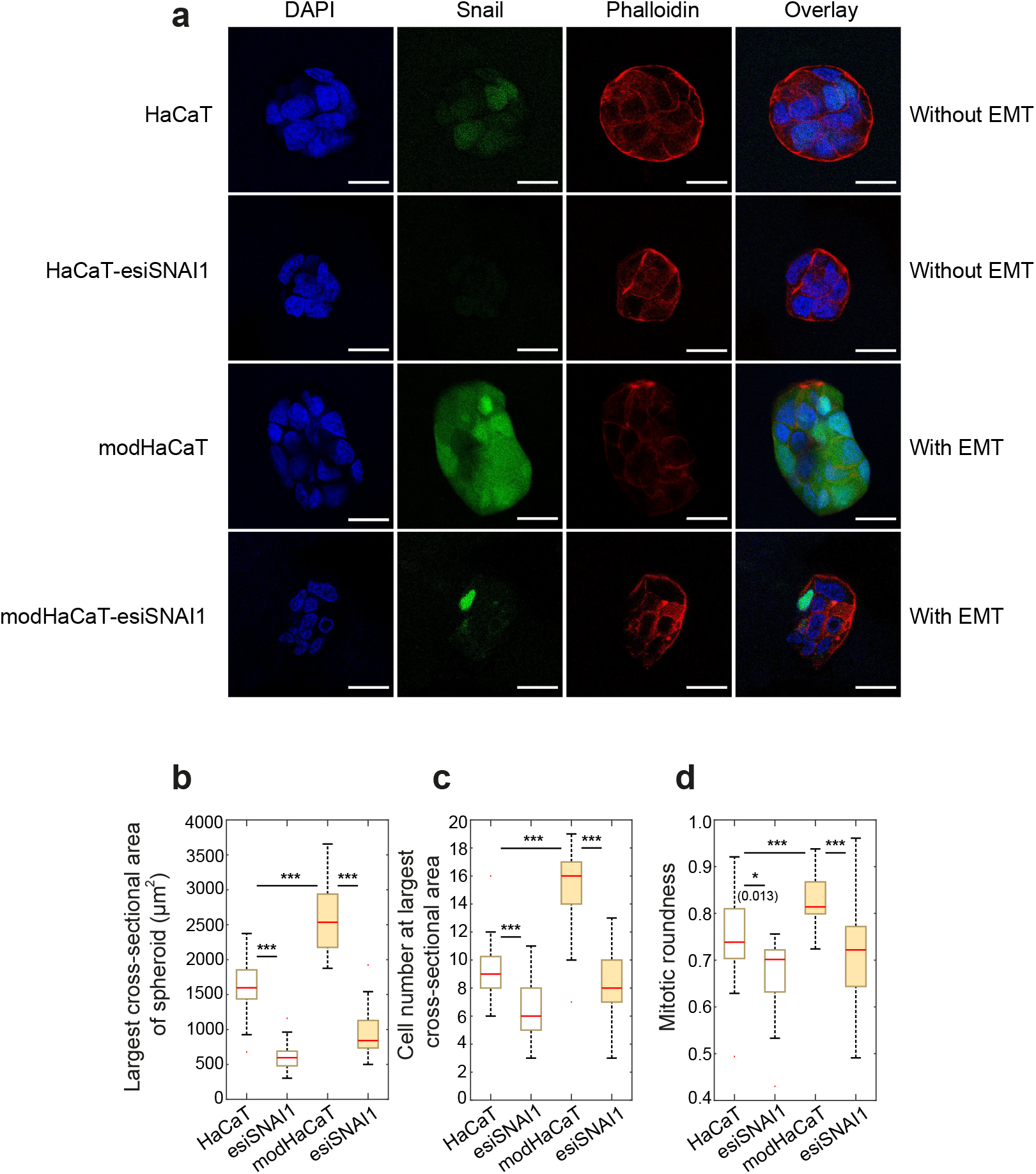
The effect of Snail knock-down on proliferation and mitotic roundness in HaCaT spheroids. a)Immunofluorescence images of cryosectioned HaCaT spheroids before and after EMT upon SNAI1 knock-down. Spheroids were stained for Snail (Green), counterstained for nuclei (DAPI, blue) and actin (Phalloidin, red). Scale bar: 20*μ*m. b-d) Spheroid size, cell number and mitotic roundness in the spheroid with and without EMT upon SNAI1 knock-down in PEG-heparin hydrogels (Young’s modulus of ≈3 kPa). b) Spheroid size was quantified by the largest cross-sectional area of the spheroid. c) Cell number was quantified as the number of cells in the maximum cross-sectional area of the spheroid (see Experimental section). Post-EMT HaCaT cells are referred to as modHaCaT. Yellow-shaded boxes indicate post-EMT conditions. Number of spheroids analyzed: (b-c): HaCaT n = 29, esiSNAI1 n= 32, modHaCaT n = 30, esiSNAI1 n= 32. Number of mitotic cells analyzed (d): HaCaT n = 20, esiSNAI1 n= 16, modHaCaT n = 35, esiSNAI1 n= 25. Measurements are representative for at least two independent experiments. n.s.: *p* > 0.05, * : *p* < 0.05, ** : *p* < 0.01, * * * : *p* < 0.001.

Overall, SNAI1 knock-down reduces spheroid proliferation and mitotic roundness. In post-EMT cells, we find that SNAI1 knock-down (over)compensates the effects of EMT induction on spheroid size, cell number and mitotic roundness leading to a diminished EMT-induced phenotype (Figure 5b-d).

## 3. Conclusion

In this work, we showed that induction of partial EMT in HaCaT skin epithelial cells causes cell-cycle-dependent mechanical changes. We assessed the actin cortex mechanics of suspended interphase cells and rounded STC-arrested mitotic cells in 2D culture for pre-EMT and post-EMT conditions. EMT resulted in the cortex becoming less contractile and softer in interphase while making the cortex more contractile and stiffer in mitosis (Figure 1c,e).

The here reported EMT-induced cell-mechanical phenotype of HaCaT cells is in accordance with previous observations of cell-mechanical changes after EMT induction for epithelial cells from breast, lung, and prostate tissue [11, 16, 41]. Overall, this suggests that cell-cycle-dependent EMT-induced changes of cell mechanics are widely conserved amongst cells of different tissue type [11, 16, 41]. Furthermore, our observations for interphase cells are in line with previous publications showing that interphase cancer cells with high metastatic potential have a softer cortex [11, 16, 35, 42–44]. Cell-mechanical changes being opposite in interphase and mitosis were also reported for Ras-induced breast epithelial cells [35].

In combination with EMT-induced cell-mechanical changes, we observed changes of cell cortex composition. Specifically, we found an EMT-induced decrease of cortical actin and myosin-II in interphase cells (Figure 2b, blue-shaded boxes) and an EMT-induced increase of cortical actin in mitotic cells (Figure 2b, yellow-shaded boxes). Furthermore, myosin-II expression was increased in post-EMT mitotic cells (Figure 2a). We conjecture that observed changes of cortex composition of actomyosin were likely caused by EMT-induced changes of cytoskeletal regulation including activity changes of Rho-GTPases [11, 45]. In addition to changes in actomyosin, we detected an increase of cortical vimentin in mitotic cells upon EMT (Figure S4d-e, Supplementary information). Vimentin was previously identified as another molecular regulator of cortical tension in the mitotic cortex [27]. Our data suggest that EMT-induced vimentin increase at the cortex also contributes to increased cortical tension post-EMT in mitosis (Figure S4a-c, Supplementary information). Overall, changes in cortex composition with regards to actin, myosin-II, and vimentin provide a plausible explanation for our observed cell-mechanical changes upon EMT.

In addition to changes in cortical mechanics and composition, we found an EMT-induced increase in cell proliferation and mitotic rounding in embedded HaCaT spheroids in 3D culture (Figure 4). Similar findings, were previously shown by us for breast epithelial cells [11]. Since mitotic rounding contributes to timely and successful cell division [46], increased mitotic roundness might be causally linked to higher proliferation rates post-EMT. Testing this hypothesis will be the object of future research.

Furthermore, we investigated the role of the transcription factor Snail on actin cortex mechanics and cortex composition before and after EMT. We found that knock-down of SNAI1 restored the pre-EMT phenotype of cortical mechanics and cortical actin and myosin-II in post-EMT cells (Figure 3). Our cell-mechanical findings on interphase cells are in accordance with previous work reporting cytoplasmic softening upon Snail overexpression in adherent MCF-7 interphase cells measured by passive intracellular bead-tracking microrheology [47].

Additionally, we reported here that knock-down of SNAI1 leads to a reduction of mitotic roundness and proliferation in HaCaT skin epithelial spheroids, in both pre- and post-EMT conditions (Figure 5b-d). Overall, in both 2D and 3D culture experiments, knock-down of SNAI1 led to a phenotype that was opposite to EMT-induced changes. The effect of SNAI1 knock-down was always stronger on the post-EMT cells compared to pre-EMT conditions (Figure 3 and Figure 5). We speculate this is due to the significant enhancement of Snail expression in post-EMT cells (Figure 5a, main text, and Figure S3a-b, Supplementary information).

In conclusion, we showed in this work that EMT in HaCaT skin epithelial cells changes cell mechanics in a cell-cycle dependent and Snail-dependent manner. Furthermore, we found that in confined HaCaT cell aggregates, serving as a benign tumor model, EMT induces a proliferation and mitotic rounding increase which depends on Snail signalling (Figure 5b-d). Interestingly, we saw, by contrast, that EMT gives rise to a proliferation decrease in 2D culture (Figure S1d, Supplementary information). Our findings suggest that EMT-induced changes of cytoskeletal regulation and cellular mechanics might play a currently under-appreciated role in EMT-induced hyperplasia of skin cells and corresponding formation of skin tumors [4].

## 4. Experimental section

### 4.1. Cell culture

The cultured cells were maintained as follows: HaCaT cells were grown in DMEM (PN:31966-021, Thermofisher) supplemented with 10% (v/v) fetal bovine (PN: 10270106, Thermofisher), 100 *μ*g/mL penicillin, 100 *μ*g/ml streptomycin (PN: 15140122, Thermofisher) at 37°C with 5% CO_2_. In HaCaT cells, EMT was induced by incubating cells in medium supplemented with 2 ng/mL TGF-*β*1 (PN:100-21, Peprotech) for 72 hours prior to the measurement. The first 24 hours of incubation were in serum-free condition [19].

### 4.2. AFM measurement of cells

#### 4.2.1. Experimental setup

To prepare mitotic cells for AFM measurements, approximately 10,000 cells were seeded in a cuboidal silicon cultivation chamber (0.56 *cm^2^* area, from cutting ibidi 12-well chamber; ibidi, Gräfelfing, Germany) that was placed in a 35 mm cell culture dish (fluorodish FD35-100, glass bottom; World Precision Instruments, Sarasota, FL) 1 day before the measurement so that a confluency of ~30% was reached on the measurement day. Mitotic arrest was induced by supplementing S-trityl-L-cysteine (Sigma-Aldrich) 2-8 hours before the measurement at a concentration of 2 *μ*M. For measurement, mitotic-arrested cells were identified by their shape. Their uncompressed diameter ranged typically from 18 to 23 *μ*m.

To prepare AFM measurements of suspended interphase cells, cell culture dishes (fluorodish FD35-100) and wedged cantilevers were plasma-cleaned for 2 minutes and then coated by incubating the dish at 37^°^C with 0.5 mg/mL poly(L-lysine)-polyethylene glycol dissolved in phosphate-buffered saline (PBS) for at least 1 hour (poly(L-lysine)(20)-g[3.5]-polyethylene glycol(2); SuSoS, Dubendorf, Switzerland) to prevent cell adhesion. Before measurements, cultured cells were detached by the addition of 0.05 trypsin-EDTA (Invitrogen). Approximately 30,000 cells in suspension were placed in the coated culture dish.

The culture medium was changed to CO_2_-independent DMEM (PN:12800-017; Invitrogen) with 4 mM NaHCO3 buffered with 20 *μ*M HEPES/NaOH (pH 7.2), for AFM experiments ~2 hours before the measurement [16, 21–23].

The experimental setup included an AFM (Nanowizard I; JPK Instruments, Carpinteria, CA) that was mounted on a Zeiss Axiovert 200M optical, wide-field microscope using a 20x objective (Plan Apochromat, NA = 0.8; Zeiss, Oberkochen, Germany) along with a CCD camera (DMK 23U445 from The Imaging Source, Charlotte, NC). Cell culture dishes were kept in a petri-dish heater (JPK Instruments) at 37°C during the experiment. Before every experiment, the spring constant of the cantilever was calibrated by thermal noise analysis (built-in software; JPK) using a correction factor of 0.817 for rectangular cantilevers [48]. The cantilevers used were tipless, 200-350 *μ*m long, 35 *μ*m wide, and 2 *μ*m thick (CSC37, tipless, no aluminum; Mikromasch, Sofia, Bulgaria). The nominal force constants of the cantilevers ranged between 0.2 and 0.4 N/m. The cantilevers were supplied with a wedge, consisting of UV curing adhesive (Norland 63; Norland Products, East Windsor, NJ) to correct for the 10^°^ tilt [49]. The measured force, piezo height, and time were output with a time resolution of 500 Hz.

#### 4.2.2. Dynamic AFM-based cell confinement

Preceding every cell compression, the AFM cantilever was lowered to the dish bottom in the vicinity of the cell until it touched the surface and then retracted to ≈14 *μ*m above the surface. Subsequently, the free cantilever was moved and placed on top of the cell. Thereupon, a bright-field image of the equatorial plane of the confined cell was recorded to evaluate the equatorial radius *R_eq_* at a defined cell height h. Cells were confined between dish bottom and cantilever wedge. Then, oscillatory height modulations of the AFM cantilever were carried out with oscillation amplitudes of 0.25 *μ*m at a frequency of 1 Hz.

During this procedure, the cell was on average kept at a normalized height *h/D* between 60 and 70%, where *D* = 2(3/(4π)V)^1/3^ and V is the estimated cell volume. Using molecular perturbation with cytoskeletal drugs, we could show in previous work that at these confinement levels, the resulting mechanical response of the cell measured in this setup is dominated by the actin cortex (see Figure 4 in [21] and Figure S7 in [11]). This is further corroborated by our observation from previous work that the smallest diameter of the ellipsoidal cell nucleus in suspended interphase cells of all cell lines under consideration is smaller than 60% of the cell diameter (see Figure S4 in [16]). Additionally, it has been shown that for a nucleus-based force response, when measuring cells in suspension with AFM, a confinement of more than 50% of the cell diameter is needed [50].

It is noteworthy that the measured cortical elastic modulus increases with cell-confinement height (up to 30% in the confinement height range of 60-70%). This phenomenon can be attributed to varying contributions of area shear and area dilation during cortical deformation, as was pointed out by our previous study [25]. Therefore, for each experiment, different conditions were measured on the same day with the same average cell-confinement height (± 6%) and the same AFM cantilever.

#### 4.2.3. Data analysis

The data analysis procedure was described in detail in an earlier work [21]. In our analysis, the force response of the cell is translated into an effective cortical tension *γ* = *F*/[*A_con_*(1/*R*_1_ + 1/*R*_2_)], where *A_con_* is the contact area between confined cell and AFM cantilever and *R*_1_ and *R*_2_ are the radii of principal curvatures of the free surface of the confined cell [16, 21, 22]. Oscillatory force and cantilever height readouts were analyzed in the following way: for every time point, effective cortical tension *γ_eff_* and surface area strain *ϵ_A_*(*t*) = (*A*(*t*) – < *A* >)/ < *A* > were estimated. An amplitude and a phase angle associated to the oscillatory time variation of effective tension γ and surface area strain are extracted by sinusoidal fits. To estimate the value of the complex elastic modulus at a distinct frequency, we determine the phase angles *φ_γ_* and *φ_ϵ_* as well as amplitudes *A_γ_* and *A_ϵ_* of active cortical tension and surface area strain, respectively. The complex elastic modulus at this frequency is then calculated as *A_γ_*/*A_ϵ_* exp(*i*(*φ_γ_* – *φ_ϵ_*)).

Statistical analyses of cortex mechanical parameters were performed in MATLAB using the commands ”boxplot” and ”ranksum” to generate boxplots and determine p-values from a Mann-Whitney U-test (two tailed), respectively.

### 4.3. Plasmids and transfection

Transfection of cells was performed transiently with plasmid DNA using Turbofectin 8.0 (PN: TF81001, Origene), according to the manufacturer’s protocol. To achieve post-EMT conditions, HaCaT cells were seeded and treated with 2 ng/mL TGF-*β*1 at day −1 in serum-starved conditions. The cells were then supplemented with serum and transfected at day 0. At day 1 the media was exchanged, removing DNA-complexes but keeping the TGF-*β*1. The Cells were then imaged at day 2. The plasmid MApple-LC-Myosin-N-7 was a gift from Michael Davidson (Addgene plasmid 54920; *http://n2t.net/addgene* : 54920; *RRID* : *Addgene*_54920). The plasmid MCherry-Actin-C-18 was a gift from Michael Davidson (Addgene plasmid 54967; *http://n2t.net/addgene* : 54967; *RRID* : *Addgene*_54967 [51]).

### 4.4. Imaging of transfected cells

The transfected cells were placed on PLL-g-PEG coated fluorodishes (FD35-100) with CO_2_-independent culture medium (described before). Cells were stained with Hoechst 33342 solution (PN:62249, Invitrogen) in order to distinguish between mitotic and interphase cells. During imaging they were maintained at 37°C using ibidi heating stage. Imaging was done using a Zeiss LSM700 confocal microscope of the CMCB light microscopy facility, incorporating a Zeiss C-Apochromat 40x/1.2 water objective. Images were taken at the equatorial diameter of each cell at the largest cross-sectional area.

### 4.5. Cortex-associated actin, myosin-II and vimentin quantification

This has been described before [11]. In short, using a MATLAB custom code, the cell boundary was identified (Figure S2b, Supplementary information shows exemplary cell, the cell boundary is marked in red). Along this cell boundary 200 radial, equidistant lines were determined by extending 1 *μ*m to the cell interior and 1 *μ*m into the exterior (Figure S2b, Supplementary information, red lines orthogonal to cell boundary, only every tenth line was plotted out of 200). The radial fluorescence profiles corresponding to these lines were averaged over all 200 lines (Figure S2c, Supplementary information, blue curve). This averaged intensity profile is then fitted by a function described in [11]. The 2D normalized cortical and cytoplasmic fluorescence intensities were obtained and used to measure the cortex-to-cytoplasm ratio in the boxplots in Figure 2b, Figure 3d main text and Figure S2d, Figure S4e Supplementary information.

### 4.6. Immunostaining and confocal imaging of cells

The suspended (interphase or STC-arrested mitotic) cells were fixed with 4% PFA/PBS for 10 minutes at room temperature, followed by a 10 min permeabilization step in 0.2% Triton X-100. The cells were then blocked for 1 hour at room temperature with 5%BSA/PBS. The cells were then treated with primary antibody of MYH9 (PN:3403T, Cell Signaling Technology) at a dilution of 1:100 or vimentin (PN:AMF-17b, Developmental Studies Hybridoma Bank, Iowa City, IA) at a dilution of 1:40 overnight at 4°C in 5%BSA/PBS. Cells were then treated with the corresponding secondary Alexa Fluor 488 conjugate at a concentration of 1:2000 in 5%BSA/PBS for 2 hours at room temperature. At the same time, cells were treated with 5 *μ*g/mL DAPI (2 minutes) and 0.2 *μ*g/mL Phalloidin-iFluor-647 (10 minutes) in 5% BSA/PBS solution. Images were taken with a Zeiss LSM700 confocal microscope of the CMCB light microscopy facility, incorporating a Zeiss C-Apochromat 40x/1.2 water objective. Images were taken at the equatorial diameter of each cell showing the largest cross-sectional area.

### 4.7. Western blotting

Protein expression in HaCaT cells before and after EMT was analyzed using Western blotting. Cells were seeded onto a 6-well plate and grown up to a confluency of 80-90% with or without EMT-inducing agents. Thereafter, cells were lysed in SDS sample/lysis buffer (62.5 mM TrisHcl pH 6.8, 2% SDS, 10% Glycerol, 50 mM DTT and 0.01%Bromophenolblue). Cell lysates were incubated for 30 minutes with the lysis buffer at 4° C. They were then boiled for 10 minutes. 20 *μ*L of the cleared lysate was then used for immunoblotting. The cleared lysates were first run on precast protein gels (PN:456-1096 or 456-1093, Bio-Rad) in MOPS SDS running buffer (B0001, Invitrogen). Subsequently, proteins were transferred to Nitrocellulose membranes (GE10600012, Sigma-Aldrich). Nitrocellulose membranes were blocked with 5% (w/v) skimmed milk powder (T145.1, Carl Roth, Karlsruhe, Germany) in TBST (20 mM/L Tris-HCl, 137 mM/L NaCl, 0.1% Tween 20 (pH 7.6)) for 1 h at room temperature followed by washing with TBST, and incubation at 4°C overnight with the corresponding primary antibody diluted 1:300 (vimentin), 1:500 (N-cadherin), 1:1000 (E-cadherin, MYH9, Snail), 1:5000 (GAPDH) or 1:50,000 (ActB) in 5% (w/v) bovine serum albumin/TBST solution. Thereupon, the blots were incubated with appropriate secondary antibodies conjugated to horseradish peroxidase, Goat anti-mouse HRP (PN: ab97023, Abcam) or Goat anti-rabbit HRP (PN: ab97051, Abcam) at 1:5000 dilution in 5% (w/v) skimmed milk powder in TBST for 1 h at room temperature. After TBST washings, specifically bound antibodies were detected using Pierce enhanced chemiluminescence substrate (ECL) (PN:32109, Invitrogen). The bands were visualized and analyzed using a CCD-based digital blot scanner, ImageQuant LAS4000 (GE Healthcare Europe, Freiburg, Germany). Primary antibodies used are as follows: E-cadherin (PN:60335-1-lg, Proteintech), N-cadherin (PN:13116T, Cell Signaling Technology), vimentin (PN:AMF-17b, Developmental Studies Hybridoma Bank, Iowa City, IA), MYH9 (PN:3403T, Cell Signaling Technology), ActB (PN:66009-1-lg, Proteintech), Snail (PN:3895, Cell Signaling Technology) and GAPDH (PN:ab9485, Abcam).

### 4.8. Preparation of spheroids in starPEG-Heparin hydrogels

As described before [11, 52], in situ assembling biohybrid multiarmed poly(ethylene glycol) (starPEG)-heparin hydrogels, prepared as previously reported [53] were used for embedding cells to grow into spheroids. In brief, cells were detached from the tissue flasks and resuspended in PBS. Subsequently, they were mixed with freshly prepared heparin-maleimide-8 solution at a density of ≈ 5 × 10^5^ cells per mL, and a final heparin-maleimide concentration of 3 mM [54]. PEG precursors were reconstituted at a concentration of 3 mM and put in an ultrasonic bath for 10-20 s in order to dissolve. 25 *μ*L of the heparin-cell suspension was mixed with equal volume of PEG solution in a pre-chilled Eppendorf tube, using a low binding pipette tip. Thereafter, a 25 *μ*L drop of PEG-heparin-cell mixture was pipetted onto a hydrophobic glass slide. After gelling, hydrogels were gently detached from the glass slide and placed in a 24-well plate supplemented with the cell culture medium. The spheroids were allowed to grow in the non-degradable hydrogel constructs for 7 days including the chemical treatment time for EMT induction, before being fixed and immunostained. Medium was exchanged every second day. After 6 days of growth, the medium was supplemented with 2 *μ*M STC to enrich mitotic cells 24 hours prior to fixation. Data are presented in Figure 4 and Figure 5.

### 4.9. Gene knock-Down through RNA interference

Transfections were done using primary esiRNA (Eupheria Biotech, Dresden, Germany) targeting the genes SNAI1 or VIM at a concentration of 25 nM, incorporating the transfection reagent Lipofectamine RNAiMax (Invitrogen). Firefly luciferase esiRNA (FLUC) was used as a negative control, while Eg5/kif11 esiRNA was used as a positive control. In all experiments, Eg5/kif11 caused mitotic arrest of more than 60-70% of the cells, showing a transfection efficiency of at least 60% in each experiment.

For AFM measurements, at day −1, 30,000 cells were seeded into a 24-well plate (NuncMicroWell Plates with Nunclon; Thermo Fisher Scientific, Waltham, MA, USA). At day 0 the transfection was done. The medium was exchanged at day 1 removing the RNAi. The transfected cells were imaged at day 2. For post-EMT conditions, the cells were kept in 2 ng/mL TGF*β*1 from day −1 to day 2 with the first 24 hours being in serum-starved conditions. For mitotic cells, ≈24 hours before measurements, cells were detached, diluted, and transferred onto a glass-bottom Petri dish (FD35-100, World Precision Instruments) with 2 *μ*M STC. For interphase cells, 2-3 hours before measurement the cells were detached and transferred to PLL-g-PEG-coated Petri dishes (see Section on AFM Measurements of Cells).

The cell volumes upon knock-down is shown in Figure S3c, Supplementary information. The data indicates an increase of cell volume upon SNAI1 knock-down in pre-EMT interphase cells (Figure S3c, upper row, white boxes, Supplementary information).

SNAI1 knock-down efficiency was also checked with immunoblotting, showing an effective reduction of Snail expression upon RNAi knockdown (Figure S3a-b, Supplementary information). Vimentin knock-down efficiency was checked with immunofluorescence, showing an effective reduction of cortical vimentin upon RNAi knock-down (Figure S4d, Supplementary information).

For spheroids, at day 5 of growth, the transfection was performed, with media exchange at day 6. The spheroids were fixed and stained at day 7. For post-EMT conditions, at day 4, 2 ng/mL of TGF-*β*1 was added in serum-starved condition. Serum was added on day 5, and TGF-*β*1 was kept until fixation. To enrich the number of mitotic cells, 2*μ*M STC was added at day 6, 24 hours before fixation, to the samples.

The SNAI1 knock-down efficiency in the spheroids was checked with immunostaining of cryosectioned slides containing spheroid slices. We observed a qualitative reduction of Snail expression upon SNAI1 knock-down, showing an effective RNAi knock-down (Figure 5a).

### 4.10. Immunostaining and confocal imaging of spheroids in hydrogel constructs

The spheroid hydrogel constructs were fixed after 7 days with 4% formaldehyde/PBS for 1 hour, followed by a 30 minutes permeabilization step in 0.2% Triton X-100. The spheroids were then stained for 4 hours with 5 *μ*g/mL DAPI and 0.2 *μ*g/mL Phalloidin-iFluor-647 in 2% BSA/PBS solution. Then hydrogels were immersed at least for 1 hour in PBS. For imaging, hydrogels were cut horizontally with a blade and were placed on a cover slip to image the interior of the hydrogel. Per hydrogel 5-10 spheroids were imaged recording a confocal z-stack with a Zeiss LSM700 confocal microscope of the CMCB light microscopy facility, using a Zeiss C-Apochromat 40x/1.2 water objective (z-step: 0.4-0.9 *μ*m).

### 4.11. Image analysis of spheroid confocal images

The software Fiji was used to characterize morphology of mitotic cells inside spheroids. Mitotic cells were identified inside a spheroid from spheroid z-stacks (red channel, F-actin, blue channel, DNA) by observing DNA structure. The largest cross-sectional area in the z-stack of each mitotic cell was determined manually by actin fluorescence, which was then used to calculate the cross-sectional shape factor roundness. Roundness was determined using Fiji by fitting an ellipse to the cross-section and calculating 4(Area)/(π(major axis)^2^), where Area is the cross-sectional area. Furthermore, spheroid size was quantified by the largest cross-sectional area of the spheroid. Data are presented in Figure 4 and Figure 5.

Statistical analysis of mitotic cell roundness in different conditions was performed in MATLAB using the commands ”boxplot” and ”ranksum” to generate boxplots and determine p-values from a Mann-Whitney U-test (two-tailed), respectively.

### 4.12. Immunofluorescence of cryosectioned spheroids

Hydrogels were fixed with 4% PFA/PBS for 1 hour at room temperature. They were then transferred to 30% sucrose/PBS solution for 1 hour at room temperature. Afterwards, they were embedded in optimum cutting temperature cryoembedding compound (OCT). They were then snap frozen in liquid nitrogen, after which they were sectioned in 10 *μ*m thick slices using Microm HM560 Cryostat (Thermofisher). The cryosectioning was performed by the BIOTEC TU Dresden histology facility. 10 minutes prior to staining, the slides were placed at room temperature and then immersed in PBS for another 10 minutes. They were then incubated with 0.5% Triton-X-100 for 1 hour at room temperature, followed by 1 hour of blocking in 5% BSA/PBS at room temperature. They were then incubated overnight at 4°C with Snail primary antibody (PN: 3895, Cell Signaling Technology) diluted at 1:200 in 1% BSA/PBS. After incubation and washing with PBS, they were incubated with corresponding secondary antibody (Alexa Fluor 488 conjugate), DAPI and Phalloidin-iFluor-647 for 3 hours at room temperature. The slides were then washed in PBS and imaged with a LSM700 Zeiss confocal microscope of the CMCB light microscopy facility incorporating a Zeiss C-Apochromat 40x/1.2 water objective. We observed that Snail expression is increased upon EMT in modHaCaT spheroids (row 1 and row 3). The images also show an efficient knock-down by the RNAi in both pre- and post-EMT spheroids. Data are shown in Figure 5a.

## Author Contributions

K.H., P.T. and A.F. performed the experiments. K.H. and E.F.-F. designed the experiments. K.H. and P.T. performed data analysis. C.W. provided reagents for hydrogel production in spheroid experiments. K.H. and E.F.-F. wrote the manuscript.

## Acknowledgements

The authors thank the CMCB light microscopy facility for excellent support. We also thank the histology facility of Biotechnology center of TU Dresden, in particular Susanne Weiche for performing the cryosectioning of the spheroids. Further, K.H., A.F. and E.F.-F. were supported by the Deutsche Forschungsgemeinschaft (DFG, German Research Foundation) under Germany’s Excellence Strategy – EXC-2068 – 390729961 – Cluster of Excellence Physics of Life of TU Dresden. Further, this project was supported by the Deutsche Forschungsgemeinschaft (DFG, German Research Foundation) by the grant FI 2260/7-1.

