## Supplementary Information for "Skin epithelial cells change their mechanics and proliferation upon Snail-mediated EMT signalling"

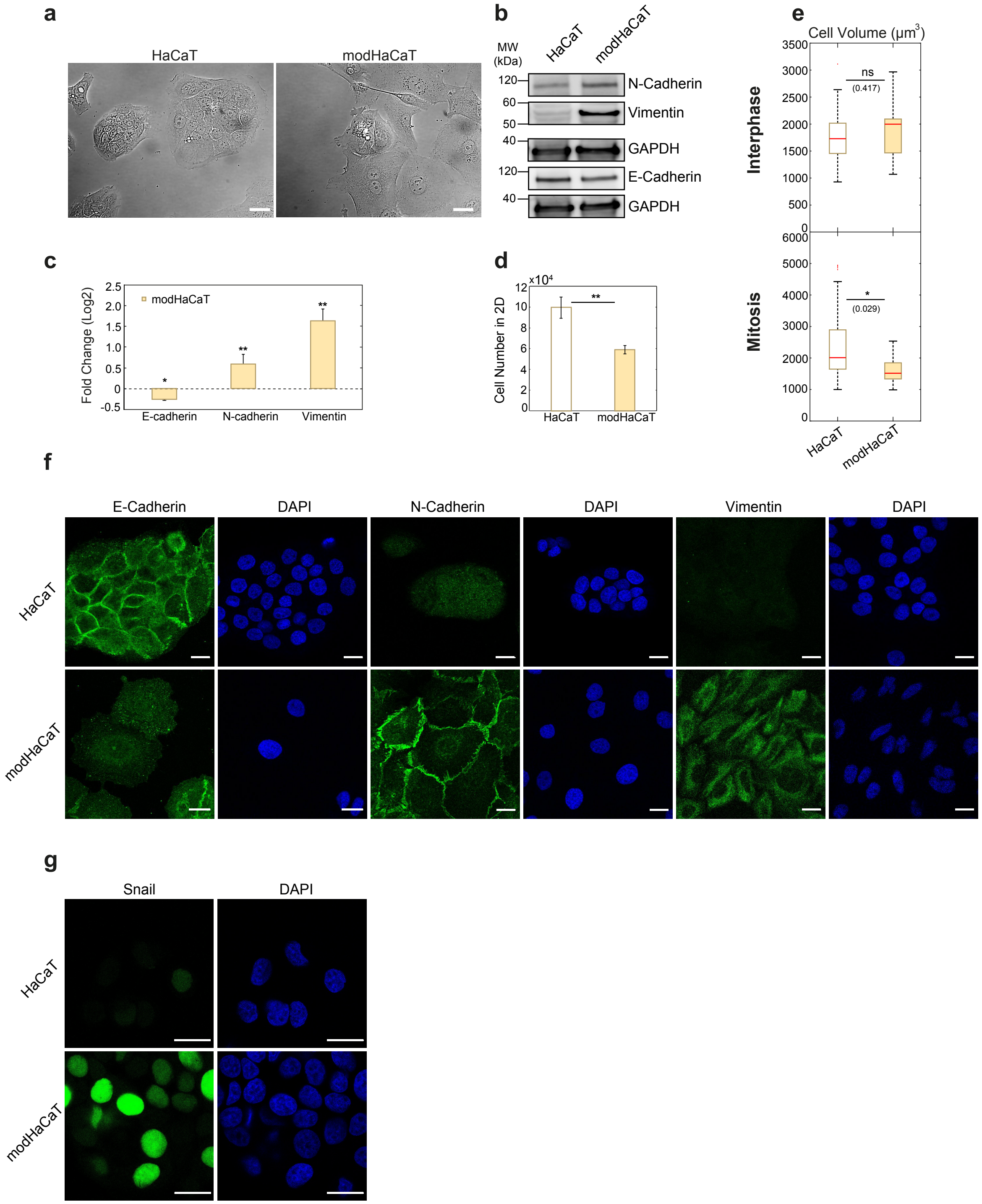


Figure S1. Pharmacological induction of EMT in HaCaT cells through TGF-$\beta$1 coincubation. a) DIC-micrograph of HaCaT (left) and EMT-induced modHaCaT (right) cells. Untreated HaCaT cells grew in clusters while EMT-transformed cells grew more isolated from each other and showed a trend towards a more spindle-shaped phenotype. Scale bar: 20 $\mu$m. b) Western blots of E-cadherin (epithelial marker), N-cadherin and vimentin (mesenchymal markers) from cell lysates before and after EMT. c) Quantification of relative changes of protein levels before and after EMT from Western blot assays (E-cadherin n=3, N-cadherin n=3 and vimentin n=3). Error bars indicate standard deviations. d) Number of cells grow in 2D culture with and without pharmacological treatment for EMT induction (see Experimental section, main text). P-values were calculated with a two-tailed student t-test. Error bars indicate standard deviations. e) Cell volumes measured for HaCaT and modHaCaT suspended interphase cells (top row) and cells in mitotic arrest (bottom row) corresponding to measurements presented in Figure 1, main text. f-g) Immunofluorescence of E-cadherin, N-cadherin, vimentin and Snail upon EMT in adherent HaCaT cells. Scale bar: 20 $\mu$m Post-EMT cells are referred to as modHaCaT. Yellow-shaded boxes indicate post-EMT conditions. Panel d: HaCaT n=5, modHaCaT n=5. Panel e: Number of cells measured: Interphase: HaCaT n = 36, modHaCaT n = 29, Mitosis: HaCaT n = 20, modHaCaT n = 17. Post-EMT cells are referred to as modHaCaT. n.s.: $p > 0.05, * : p < 0.05$, $** : p < 0.01$, $*** : p < 0.001$.

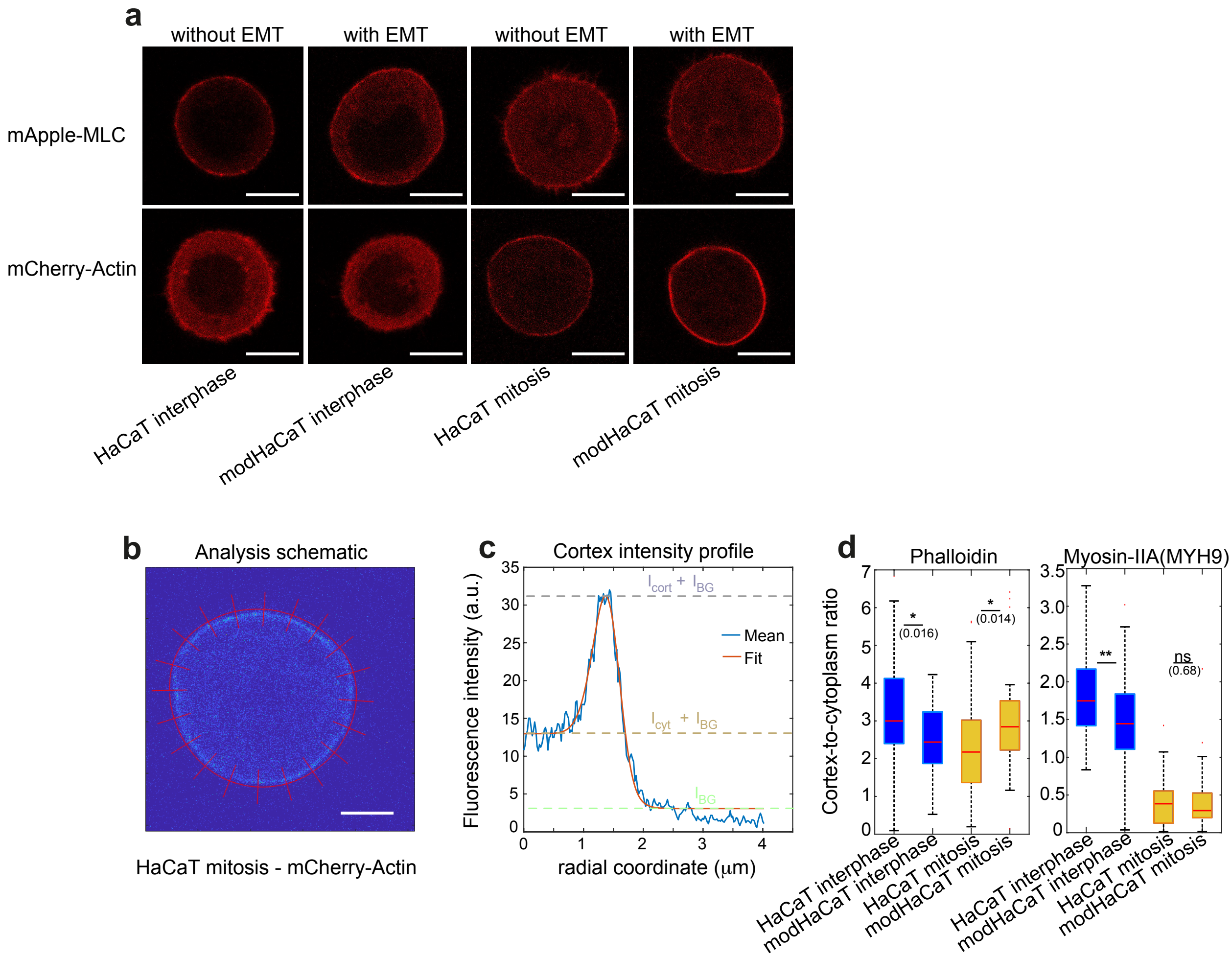


Figure S2. Actin and myosin-II changes in the cortex of skin epithelial cells upon EMT. a) Representative confocal images of suspended interphase cells and STC-arrested mitotic cells expressing mCherry-ACTB or mApple-Myl9, with and without EMT. Scale bar: 10 $\mu$m. b) Exemplary picture of myosin-II fluorescence profile of the equatorial cross-section of a mitotic cell including elements of image analysis. Scale bar: 10 $\mu$m. c) Mean radial fluorescence intensity profile (blue curve) along radial lines shown in panel b. The fitted intensity profile, $I_{sm}(r,p)$, (see Experimental section, main text) is shown in orange. d) Ratio of cortical versus cytoplasmic F-actin and myosin-IIA intensity (labelled with Phalloidin and MYH9 antibody) in fixed suspended interphase cells and STC-arrested mitotic cells, pre- and post-EMT (see Experimental section, main text). Post-EMT cells are referred to as modHaCaT. Blue-shaded boxes indicate interphase cells while yellow-shaded boxes indicate mitotic cells in (d). Number of cells measured: Panel d: Phalloidin: HaCaT interphase n=47, modHaCaT interphase n=49, HaCaT mitotic n=46, modHaCaT mitotic n=39, MYH9: HaCaT interphase n=48, modHaCaT interphase n=47, HaCaT mitotic n=47, modHaCaT mitotic n=48. Measurements are representative for two independent experiments. n.s.: $p > 0.05, * : p < 0.05$, $** : p < 0.01$, $*** : p < 0.001$.

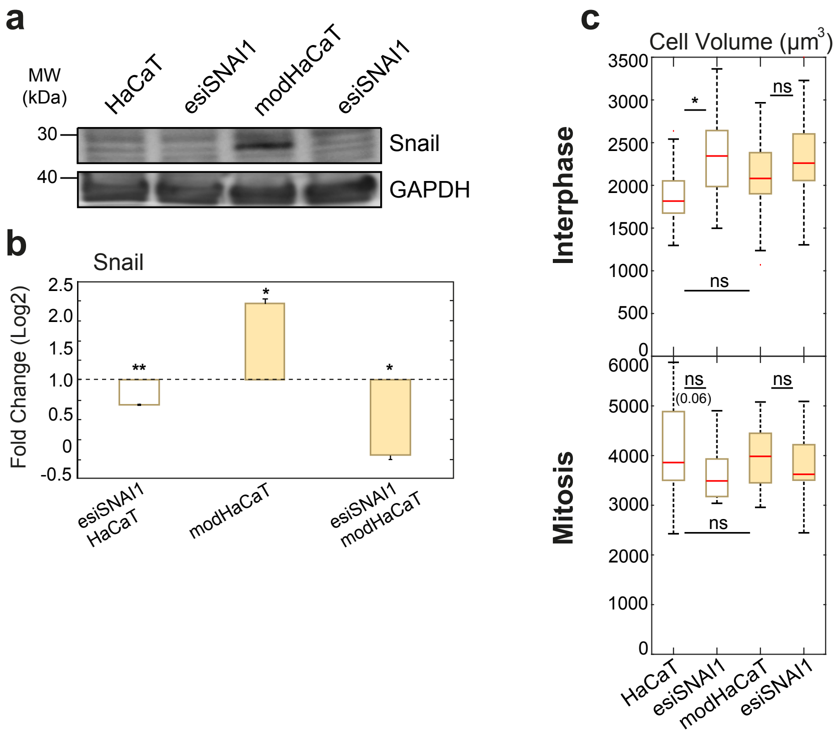


Figure S3. Relative changes of Snail expression upon EMT. a,b) Western blots and quantification showing Snail abundance in lysates of HaCaT and modHaCaT (in asynchronous cell populations) in control conditions and upon SNAI1 knock-down (n=2). Knock-down was achieved through RNA interference. Quantification was done against GAPDH. The Western blots show successful expression changes of the target protein upon knock- down or pharmacological treatments. c) Cell volume measured for HaCaT and modHaCaT suspended interphase cells (top row) and cells in mitotic arrest (bottom row) upon SNAI1 knock-down corresponding to measurements presented in Figure 3, main text. Post-EMT cells are referred to as modHaCaT. Yellow-shaded boxes indicate post-EMT conditions. Number of cells measured: Interphase: HaCaT n = 27, esiSNAI1 n=28, modHaCaT n = 27, esiSNAI1 n=29, Mitosis: HaCaT n = 22, esiSNAI1 n=20, modHaCaT n = 20, esiSNAI1 n=19. Measurements are representative for two independent experiments. n.s.: $p > 0.05, * : p < 0.05$, $** : p < 0.01$, $*** : p < 0.001$.

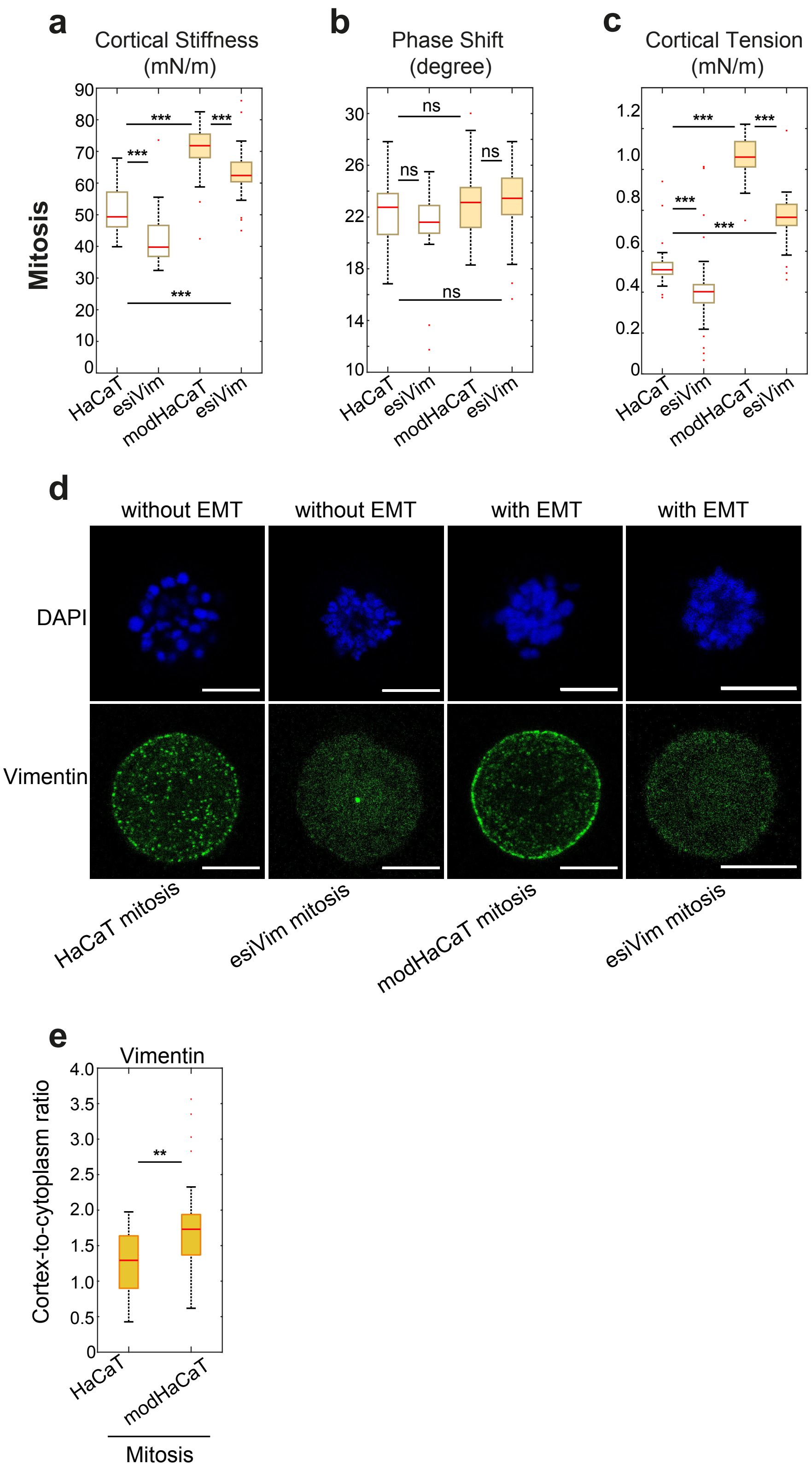


Figure S4. Effect of vimentin knock-down on cortex mechanics in STC-arrested mitotic HaCaT cells with and without EMT. a) Cortical stiffness, b) phase shift, and c) cortical tension measured by dynamic AFM confinement. d) Representative images of stained STC-arrested mitotic HaCaT cells: vimentin immunofluorescence (green) and DAPI (blue), with and without EMT. Scale bar: 10 $\mu$m. e) Ratio of cortical versus cytoplasmic vimentin in STC-arrested mitotic cells upon EMT. Post-EMT HaCaT cells are referred to as modHaCaT. Yellow-shaded boxes in (a-c) indicate post-EMT conditions. Yellow-shaded boxes in (d) indicate mitotic cells. Number of cells measured in (a-c): HaCaT n = 28, esiVim n = 27, modHaCaT n = 26, esiVim n = 26. (e): HaCaT n = 36, modHaCaT n = 37. Measurements are representative for at least two independent experiments. n.s.: $p > 0.05, * : p < 0.05, ** : p < 0.01, *** : p < 0.001$.
